# Adaptation of Intracortical Signaling Concurs with Enhanced Encoding Efficiency

**DOI:** 10.1101/320226

**Authors:** Jacob A. Westerberg, Michele A. Cox, Kacie Dougherty, Alexander Maier

## Abstract

Stimulus repetitions improve performance despite decreased brain responses, suggesting that the brain is more efficient when processing familiar stimuli. Previous work demonstrated that stimulus repetition enhances encoding efficiency in primary visual cortex (V1) by increasing synchrony and sharpening the orientation tuning of neurons. Here we show that these adaptive changes are supported by an altered flow of sensory activation across the V1 laminar microcircuit. Using a repeating stimulus sequence, we recorded laminar responses in V1 of two fixating monkeys. We found repetition-related response reductions that were most pronounced outside V1 layers that receive the main retinogeniculate input. This repetition-induced suppression was robust to alternating stimuli between the eyes, in line with the notion that repetition suppression is predominantly of cortical origin. Congruent with earlier reports, we found that V1 adaptation to repeating stimuli is accompanied by sharpened neural tuning as well as increased neural synchrony. Current source density (CSD) analysis, which provides an estimate of net synaptic activation, revealed that the responses to repeated stimuli were most profoundly affected within layers that harbor the bulk of cortico-cortical connections. Together, these results suggest that stimulus repetition induces an altered state of intracortical processing resulting in enhanced encoding efficiency of sensory stimuli.

---

The visual system relies on prior experiences to derive behaviorally relevant interpretations from sensory information (Helmholtz 1866). One well-studied, experience-related brain process is the reduction of neuronal responses to repetitions of the same or similar stimuli (Barron et al. 2016). On the level of single neurons, this kind of adaptation has been referred to as “repetition suppression” (Baylis and Rolls 1987; Miller et al. 1991; Riches et al. 1991; Li et al. 1993; Miller et al. 1993; Kobatake and Tanaka 1994; Miller and Desimone 1994; Sobotka and Ringo 1994, 1996; Sawamura et al. 2006; McMahon and Olson 2007; Liu et al. 2009; Kaliukhovich and Vogels 2014). Repetition suppression of visual spiking responses occurs even if the repeating stimuli are not identical (Liu et al. 2009), but gradually falls off in magnitude with decreasing stimulus similarity (Sobotka and Ringo 1994). Repetition suppression thus consists of both a stimulus-specific as well as a generalized adaptation component (Kohn 2007) and occurs under a wide variety of circumstances where stimuli are repeatedly encountered (Riches et al. 1991; Sawamura et al. 2006).

Interestingly, observers become both faster and more accurate at responding to visual stimuli when they have been previously exposed to the same or similar stimuli (Tulving and Schacter 1990). Given the reduced neuronal response to repetitions that have been observed under similar stimulation conditions, these behavioral improvements need to be mediated by mechanisms other than response magnitude. Previous work identified at least two spike rate-independent processes that underlie repetition-induced enhancement of encoding efficiency (Gotts et al. 2012). Specifically, work in humans (Gilbert et al. 2010) and monkeys (Hansen and Dragoi 2011; Brunet et al. 2014) found that repetition increases synchrony as well as neuronal selectivity (tuning).

Here, we demonstrate that these previously uncovered mechanisms of enhanced encoding efficiency are supported by systematic changes in intracortical processing within the laminar microcircuitry of primary visual cortex (V1). Using linear multielectrode arrays in two fixating monkeys, we recorded neuronal responses to repeated stimulation across all V1 laminae. Laminar response activity profiles are informative as the granular input layer 4C receives the bulk of sensory input, while the layers above and below harbor the bulk of intracortical processing (Glickstein et al. 1967; Garey and Powell 1971; Rockland and Pandya 1979; Rockland and Virga 1989; Felleman and Van Essen 1991; Mignard and Malpeli 1991; Anderson and Martin 2009; Markov et al. 2014). We found that stimulus repetition produces reductions in firing rates across all V1 layers. Further examination revealed this suppression was largest in supragranular layers (layers 2/3) where the bulk of intracortical processing occurs. In line with earlier findings (Hansen and Dragoi 2011), we found that both the strength of orientation tuning of V1 neurons as well as V1 spike-field coherence increased with repetitive stimulation despite reduced spiking. Current source density analysis revealed that the main reduction of synaptic drive underlying repetition suppression was also located in supragranular layers. Finally, stimulating one eye initially and testing for repetition suppression in the other revealed that the majority of V1 neurons show similar response suppression under these conditions. These results combined suggest that repetition-related changes in V1 sensory processing are predominantly based on adaptation of intracortical signaling in the supragranular layers.

## Methods

### Animal Care and Surgical Procedures

Two adult monkeys (*Macaca radiata*), one female (monkey I34) and one male (monkey E48) were used in this study. All procedures were in compliance with regulations set by the Association for the Assessment and Accreditation of Laboratory Animal Care (AAALAC), approved by Vanderbilt University’s Institutional Animal Care and Use Committee, and followed National Institutes of Health guidelines. In a series of surgeries, each monkey was implanted with a custom-designed MRI-compatible head post and a plastic recording chamber (Crist Instrument Co., Inc.) situated over striate cortex.

All surgeries were performed under sterile surgical conditions using isoflurane anesthesia (1.5–2.0%), following induction with an intramuscular injection of ketamine hydrochloride (10mg/kg). Vital signs, including blood pressure, heart rate, SpO_2_, CO_2_, respiratory rate, and body temperature, were monitored continuously. During surgery, the head post and recording chamber were attached to the skull using transcranial ceramic screws (Thomas RECORDING GmbH), and self-curing denture acrylic (Lang Dental Manufacturing Inc.). Further, a craniotomy was performed over the perifoveal visual field representation of primary visual cortex concurrent with the position of the recording chamber. Each monkey was given analgesics and antibiotics for post-surgical care.

### Behavioral Task and Visual Stimulation

Monkeys passively fixated within a one-degree radius around a central fixation dot. All stimuli were presented with a 20” CRT monitor (Diamond Plus 2020u, Mitsubishi Electric Inc.) operating at 60 or 85Hz. We used a custom mirror stereoscope so that stimuli could be presented to either eye using either the left or the ride side of the monitor. Stimuli were generated using MonkeyLogic (Asaad and Eskandar 2008) for MATLAB (R2012, R2014a, The Mathworks, Inc.). After a 300ms fixation period, a sequence of five static, sinusoidal stimuli of equivalent size, spatial frequency, and phase with variable orientation were presented while the monkey maintained fixation. Exact values for the stimulus parameters as well as spatial location of the stimuli were selected dependent upon the response properties of the neurons recorded in each session, as evaluated by listening to the auditory multi-unit activity (MUA) to a variety of stimuli across several randomly chosen channels (0.5-3cyc./deg. at 2-7dva eccentricity). In other words, we selected stimulus parameters that we deemed to evoke the largest overall response, but none of these parameters were optimized for individual neurons. In the ‘short’ stimulation duration condition, each stimulus in the sequence was presented for 200ms. In the ‘long’ stimulation duration condition, each stimulus was presented for 500ms. Both conditions featured a 200ms inter-stimulus interval (ISI) between each stimulus. At 200ms following the last stimulus of each sequence, the monkey was relieved from the constraint of fixation and received a juice reward. If at any point during the trial (stimulus sequence) the monkey broke fixation or blinked, the trial was aborted and the monkey experienced a short (1-5s) timeout before starting the next trial.

### Neurophysiological Procedure

Broadband (0.5Hz-12.207kHz) intracranial voltage fluctuations were recorded inside an electromagnetic radio frequency-shielded booth with an acute laminar multielectrode array. Voltages were amplified, filtered, and digitized using the 128-channel Cerebus™ Neural Signal Processing System (Blackrock Microsystems LLC). Two different laminar probes were used (U-Probe, Plexon, Inc.; Vector Array™, NeuroNexus). The laminar probes consisted of either 24 or 32 active microelectrodes, linearly spaced 0.1mm apart, with impedances ranging 0.2-0.8MΩ at 1kHz. The probes were connected to the Cerebus™ amplifier using an analog headstage (Blackrock Microsystems LLC). Each recording session, one or two laminar probes were introduced into dorsal V1 through the intact dura mater using a chamber-mounted microdrive (a custom-designed modification of a Narishige International Inc. Micromanipulator) and adjusted in the z-plane until the majority of microelectrode contacts spanned the entire cortical thickness, from the subdural space to the white matter. Gaze position was recorded at 1kHz (NIDAQ PCI-6229, National Instruments) using an infrared-light sensitive camera and commercially available eye tracking software (Eye Link II, SR Research Ltd.; iView, SensoMotoric Instruments). All trials where the animal’s gaze left the central one-degree radius around the fixation spot were excluded from analysis. Local field potentials (LFP) were extracted from the broadband signal using a low-pass Butterworth filter with a cutoff frequency of 500Hz. Single-unit spiking activity was sorted offline using the KiloSort toolbox (Pachitariu et al. 2016) for MATLAB (R2016b). KiloSort is specifically well adapted to linear electrode arrays such as the ones used in this study as this software automatically detects any coincident spikes on neighboring channels and uses these signals to eliminate double counting of the same spikes if they are picked up by more than one electrode channel. Visual inspection of the mean and variance of the isolated spike waveforms suggests that KiloSort spike isolation is on par with or even surpasses conventional cluster-based spike sorters (see Fig. 1C and Pachitariu et al. 2016). We used the KiloSort default parameters for sorting and cluster merging.

**Figure 1.**
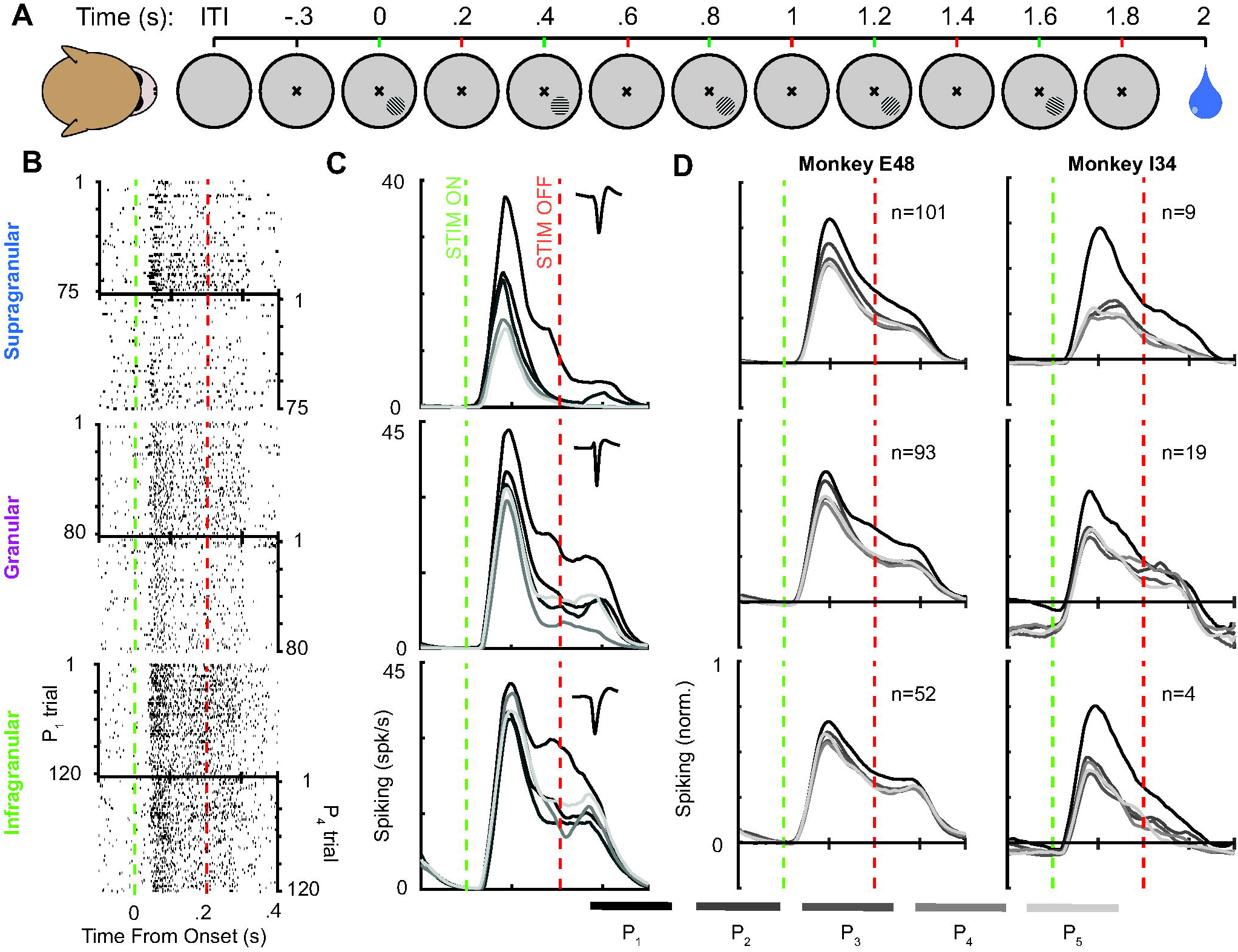
(A) Passive fixation paradigm employing repetitive visual stimulation to receptive field of neurons under study. (B) Raster plots of representative single unit responses for each of the three main laminar compartments (key left) for the first presentation (top) and fourth presentation of each sequence (bottom). Green vertical line indicates the onset of visual stimulus and red, the offset. (C) Spike density functions for the same responses. The response to each presentation is denoted by color, with black indicating first and the lightest gray indicating the fifth presentation (key beneath C, D). Insets represent the mean spiking waveforms for each respective unit. (D) Median spike density functions for each monkey across all normalized single unit responses per laminar compartment.

### Laminar Alignment

CSD analyses of visual responses to brief visual stimulation have been shown to reliably indicate the location of the primary geniculate input to V1 (to granular layer 4C, or L4) as a distinct current sink that is thought to reflect the combined excitatory post-synaptic potentials of the initial retinogeniculate volley of activation (Mitzdorf and Singer 1979; Schroeder et al. 1998). To derive CSD from the LFP, we computed an estimate of the second spatial derivative appropriate for multiple contact points using the following formula (Nicholson and Freeman 1975):

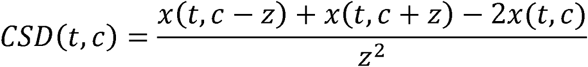

where *x* is the extracellular voltage at time t measured at an electrode contact at position *c*, and *z* is the inter-contact distance of the electrode array. The resulting CSD from the formula above was multiplied by 0.4S/mm as an estimate of the electric conductivity of cortex (Logothetis et al. 2007) to obtain current per unit volume. We eliminated channels at the top and the bottom of the electrode array that were void of any MUA visual response (see Cox et al. 2017 for details). For display purposes, two-dimensional representations of CSD as a function of time and space were created by interpolating CSD between adjacent electrode contacts, followed by smoothing with a two-dimensional Gaussian filter (*s* = 0.1mm and 15ms) (Pettersen et al. 2006). Microelectrode contacts were determined to be located in the granular layer based on the initial sink of a flash-evoked CSD (see Maier et al. 2010; Maier 2013; Dougherty et al. 2015; Ninomiya et al. 2015; Cox et al. 2017 for details). Infragranular laminar compartment locations were determined by the position of the initial sink in response to a flashed stimulus and corroborated by additional neurophysiological criteria (Maier et al. 2010), such as characteristic patterns of LFP power spectral density (van Kerkoerle et al. 2014; Bastos et al. 2018), signal correlations of the LFP between all channel combinations, and the latency (Self et al. 2013) of stimulus-evoked MUA responses (Supplementary Fig. 1). The supragranular to granular boundary was set to 0.5mm above the granular to infragranular boundary.

### Receptive Field Mapping

Once satisfactory electrode placement was achieved, we used a reverse correlation-like technique to map the receptive fields of the single units under study. We first estimated the receptive field location from the audible MUA response to bars and gratings that were moved across the screen while the animals fixated for juice reward. We then had the animals fixate while a series of circular static random noise patches were displayed at pseudorandomized locations within a pre-determined virtual grid that covered the estimated receptive field. Up to five stimuli were shown per trial, for 200ms each with 200ms blank periods interleaved. Stimulus size and grid spread varied depending on receptive field estimates, with each recording session typically including an initial “coarse” followed by a “fine” mapping phase of decreasing grid size. We used the resulting neurophysiological data to compute retinotopic 3D Receptive Field Matrices (RFMs) (Cox et al. 2013) to derive spatial maps of neuronal responses as a function of visual space. Briefly, for each stimulus presentation, the evoked neuronal response (either MUA or SUA from each microelectrode contact) was averaged over time and converted to z-score to produce a single scalar value for each trial. The portion of virtual grid corresponding to the stimulus location was then filled with that value, producing a three-dimensional matrix consisting of one dimension for vertical visual space, one dimension for horizontal visual space, and one dimension for stimulus presentations. The third dimension was then collapsed via averaging, producing a spatial map of each unit’s retinotopic sensitivity. Sessions where receptive fields did not overlap across all electrode channels were eliminated from analysis.

### Neurophysiological Analysis and Statistical Testing

All analyses of single-unit activity were performed using custom written scripts in MATLAB. Trials with exceeding electrical noise or motion artifacts were excluded using the generalized extreme studentized deviate (ESD) correction (Rosner 1983) at the unit level by computing:

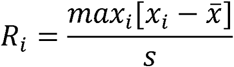

and

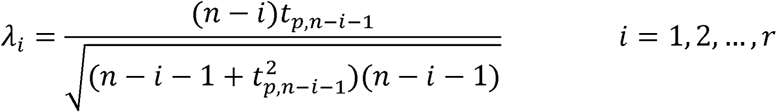

where the number of outliers for a given unit is the largest *i* such that *R_i_* is larger than λ*_i_*, *n* is the number of trials, *s* is the standard deviation of the maximum values of each trial, *x* is the maximum value of the trial, *t_p,v_* is the value of the t distribution with *v* degrees of freedom *α* = 0.05, and:

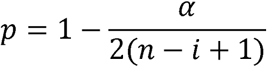

Spiking data was convolved using a Poisson distribution resembling a postsynaptic potential (Thompson et al. 1996), where the spike rate (*R*) is computed at time (*t*) where:

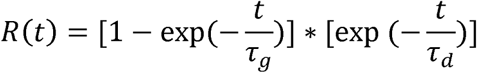

such that *τ_g_* and *τ_d_* are the time constants for growth and decay, respectively. Data from previous studies suggest values of 1 and 20 as valid for *τ_g_* and *τ_d_*, respectively (Sayer et al. 1990; Mason et al. 1991; Kim and Connors 1993; Thomson et al. 1993a, 1993b).

For investigation of within-unit conditional changes, the convolved spiking data was analyzed using a receiver operating characteristic (ROC) (Green and Swets 1966). Conditional differences were quantified by taking the average spiking rate of each unit between 40ms post-stimulus onset to stimulus offset for each repetition. These average values then served as the parameter *λ* to generate Poisson processes across values *k* of the distribution:

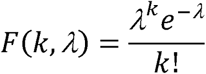

For each condition, values from the Poisson distributions were used to generate ROC curves by taking a sample of the Poisson data that matched the minimum number of trials for the compared conditions. The area under the curve was tested against a random distribution by bootstrapping 1000 times to determine whether the difference between the conditions was significant at *α =* 0.05.

Units with no significant response to visual stimulation, determined by performing a two-sample t-test (*α* = 0.05) on the mean baseline activity of each trial and the mean activity during the epoch of visual stimulation (50ms post-stimulus onset to stimulus offset), were excluded from analysis. Additionally, units whose average maximum firing rate in response to visual stimulation did not exceed five spikes per second were excluded from analysis.

SUA was made comparable across individual units with differing average spike rates by normalizing their responses at the unit level:

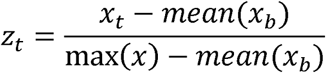

where *x_i_* is the averaged data for all stimulus orientations in each set at time *t* and *x_b_* is the averaged data restricted to the baseline epoch. The baseline epoch was defined as the 100ms interval of passive fixation leading to the onset of the stimulus. *z_t_* is the resultant normalized value at time point *t*.

Statistical tests were performed at the unit level using two-tailed, two-sample t-tests (*α* = 0.05) and adjusted using the Benjamini-Hochberg procedure for false discovery rate at (*FDR* = 0.05) (Benjamini and Hochberg 1995).

To determine the latency of neuronal responses, each unit was tested by computing two-sample t-tests (*α* = 0.05) at each time point against the baseline (prestimulation) epoch on each trial. The point at which the response deviated significantly from the baseline for five sequential samples was found to be the latency of that response. The mean latency was taken for each unit. The mean of these values for all units was taken to obtain the population average.

### Tests for Ocularity and Orientation Tuning

We used a custom-designed mirror stereoscope (see Cox et al. 2017) to evoke neuronal responses specific to each eye. For any given stimulus presentation, the stimulus could be presented to either the left eye, the right eye, or to both eyes simultaneously. We used the neuronal responses to monocular stimulation to compute an ocularity index for each unit. Similar to recent work on ocular dominance in V1 (Zhang et al. 2005; Romero et al. 2007), we computed a quantitative ocularity index (*OCI*) using a Michelson contrast (Michelson 1927) to compare single unit responses between stimulus presentations to the left and right eyes:

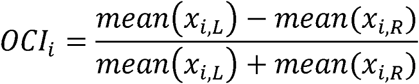

where the ocularity index for unit *i* (*OCI_i_*) was defined as the difference between the mean response *x* for unit *i* on trials where the stimulus was monocularly presented to the left eye (*x_i,L_*) and when the stimulus was presented monocularly to the right eye (*x_i,R_*) divided by the sum of those two responses.

To derive the average stimulus responses, we computed the mean spike rate between 40ms post-stimulus onset to stimulus offset. Units with a Michelson contrast above 0.6 were deemed “monocular” units while units below 0.4 were defined as “binocular” units.

We also determined each unit’s orientation preference. Specifically, activity from 40ms post-stimulus onset to stimulus offset was used to derive average responses to each stimulus type. Grating orientations were grouped into 9 bins that encompassed 20° of angle. In order to determine whether orientation preference for a given unit was significant, we performed a one-way ANOVA (*p* < 0.05) between these binned responses. In all cases, the stimulus orientation with the greatest average response was defined as a unit’s preferred orientation.

We determined repetition-dependent changes in orientation tuning by computing the tuning curves for each neuron with significant orientation, followed by normalization between zero and one. This process was performed for each stimulus repetition independently, providing normalized tuning curves for the initial stimulus presentation as well as for all ensuing repetitions. By subtracting between these tuning curves, we measured the repetition-induced change in tuning for each of these neurons.

Fitting the representative unit’s orientation tuning curve was done in several steps. First, the unit was found to have significant orientation tuning following a one-way ANOVA where at least one orientation had a significantly different response magnitude (*p* < 0.05). Following that, the preferred orientation was found by determining the stimulus orientation which evoked the greatest response. Stimulus responses to grating orientations with identical absolute differences relative to the preferred orientation were then binned. These oblique values were then duplicated and mirrored and a first-order Gaussian fit was performed.

We found that both ocular preference as well as the peak of orientation tuning was largely shared across neurons for each penetration, as expected given V1’s anatomical makeup of ocular dominance columns and orientation columns. Comprehensive quantitative assessment and statistical description of these data is beyond the scope of the current study and will be the focus of an upcoming paper.

### Local Field Potentials and Spike-Field Coherence

We used the Chronux toolbox for MATLAB (Bokil et al. 2010) for all LFP analyses. Power spectral densities were computed for the 160ms window that lasted from 40ms post stimulus onset to stimulus offset. For the baseline condition, we used a window of 160ms prior to the first stimulus onset of each trial. For spike-field coherence (SFC) analysis, spiking was tested against the continuously-sampled LFP recorded at the same electrode channel. This choice was for the following reasons: 1) Our goal was to determine SFC as a function of laminar compartment. We thus wanted to make sure that both the spikes and the LFP were not sampled across laminar boundaries. 2) Spikes sometimes span multiple electrode contacts on linear arrays, so choosing neighboring contacts for spikes and LFP, respectively, does not effectively dissociate LFP and spiking waveforms. 3) We compare SFC between conditions, so potential contamination by spikes should be similar across comparisons, and 4) initial work on SFC was performed using the same technique and yielded informative results (Fries et al. 2001).

### Anatomical MRI

Animals were anesthetized using the same procedure as outlined under Animal Care and Surgical Procedures. Anesthetized animals were then placed inside a Philips Achieva 7T MRI scanner (Philips) at the Vanderbilt University Institute of Imaging Science and remained anesthetized throughout the duration of the scan. Vital signs were monitored continuously. T1-weighted 3D MPRAGE scans were acquired with a 32-channel head coil equipped for SENSE imaging. Images were acquired using a 0.5mm isotropic voxel resolution with the following parameters: repetition time (TR) 5s, echo time (TE) 2.5ms, flip angle 7°.

## Results

Adaptation on the level of neural circuits is an area of active research (Kar and Krekelberg 2016; Quiroga et al. 2016). Here we devised a stimulation sequence consisting of five stimulus repetitions to examine the effect of adaptation on the cortical laminar microcircuit. Each stimulus was a static grating of fixed spatial position, contrast, phase, size, and spatial frequency. To determine repetition-induced changes in orientation tuning (see below), we randomized stimulus orientation (see Methods). Each grating was presented for 200ms, followed by a 200ms ISI (Fig. 1A). Two monkeys performed this task (n=48 sessions in monkey E48, n=13 in I34). Following successful cortical penetration with the linear multielectrode array and careful laminar alignment (see Methods and Supplementary Fig. 1), we recorded V1 spiking responses across all layers. The five successively presented visual stimuli always spanned the receptive fields of the recorded neurons.

### Spiking Suppression During Repetitive Visual Stimulation

Across all recording sessions in both animals, we extracted 278 single units from the broadband signal using an unsupervised machine learning algorithm (Methods) (Pachitariu et al. 2016). We sampled units from the upper (supragranular n=110), middle (granular layer 4C n=112) and lower (infragranular n=56) layers, which allowed us to study the effects of adaptation on each of these main laminar compartments.

We first examined laminar single-unit activity (SUA) across consecutive stimulus presentations (Figs. 1B-D, 2), irrespective of stimulus orientation or stimulated eye (see below for a detailed analysis of each of these parameters). This comparison revealed a reduction of response magnitude to repeated stimuli. This response reduction affected both the transient as well as the sustained phase of the visual response in all layers (Figs. 1B-D, 2A, *see* below for statistics). The largest repetition-induced decrease in spiking occurred immediately following the first stimulus repetition, with smaller spike rate decrements for subsequent presentations (Figs. 1B-D, 2A, see also Supplementary Figs. 2, 3, 4).

**Figure 2.**
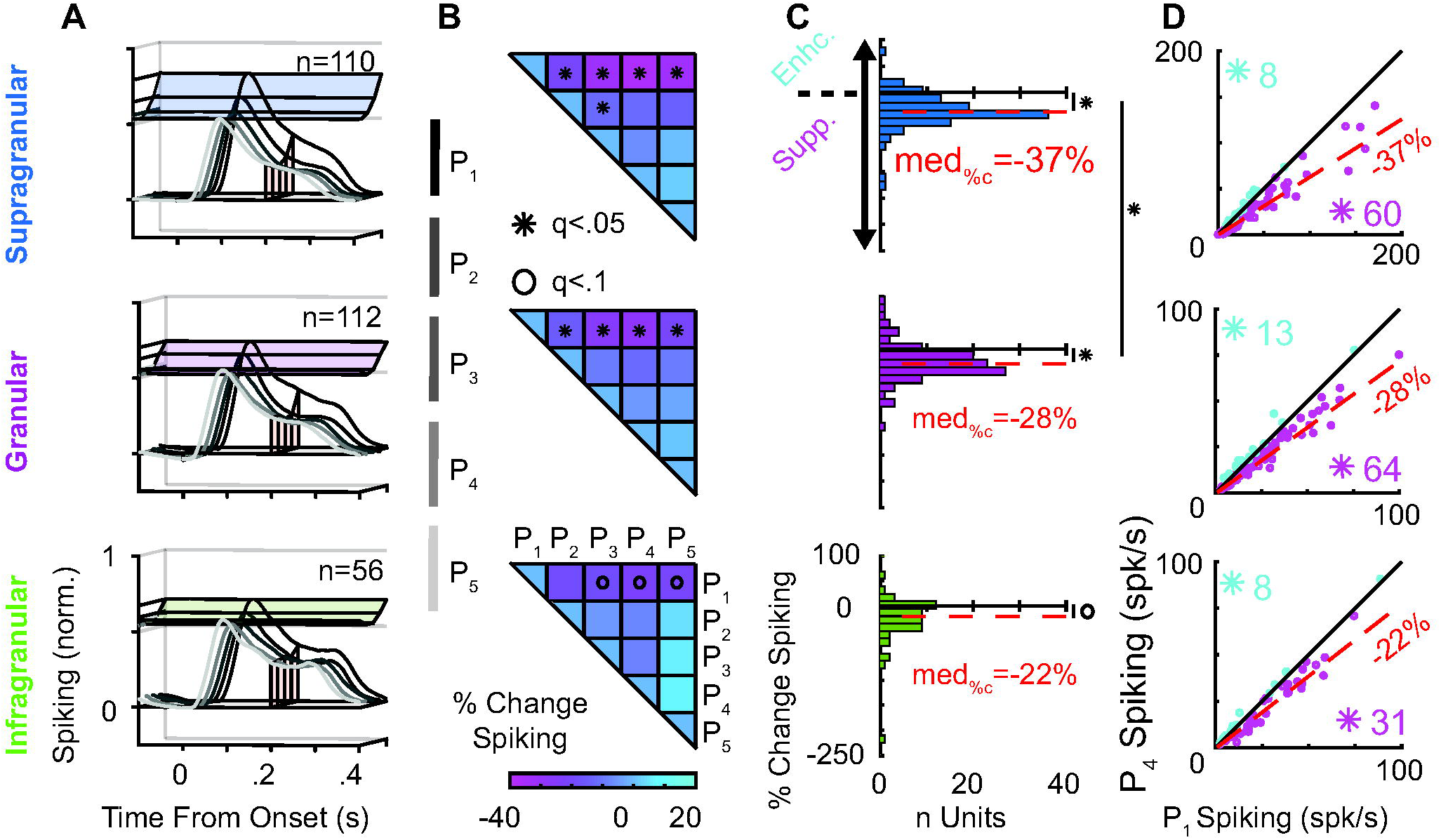
Laminar profile of V1 repetition-induced spiking suppression. (A) 3-Dimensional representation of median spike density functions for each laminar compartment across all units in both monkeys. Responses to subsequent repetitions are color coded in gray scale (legend right). Colored planes represent response maxima between subsequent presentations. (B) Change in population response between each subsequent stimulus presentation. Color represents percent change of evoked spiking. Significant differences are indicated with an asterisk (see Methods). (C) Distribution of single unit response differences between first and fourth presentations in percent change. Values below 0 denote response suppression. Median response is indicated by a red dotted line. (D) Significance of repetition-induced response differences as evaluated by a Poisson spike train analysis (see Methods). Each single unit is represented by a dot in a scatter plot where the median spiking response to the first presentation is compared to the fourth. Magenta numbers provide the number of units showing significant suppression in each compartment. Cyan numbers denote single units with significant response facilitation.

Collapsing the population average responses across time, we found that all stimulus repetitions evoked significantly smaller responses than the initial presentation in both granular layer 4C and supragranular layers, with suppression in the infragranular layers approaching significance (Fig. 2B; two-sample t-test, corrected with FDR, *α* = 0.05). Comparing population responses for presentations two through four using the same technique, we found significant differences between only the second and third presentations in the supragranular layers (Fig. 2B, top). The magnitude of this repetition suppression was largely independent of baseline or maximum firing rates (Supplementary Fig. 5), indicating that laminar position, rather than spiking characteristics, drove the effect.

As a next step, we compared each neuron’s response between the initial and the fourth presentations (since the fourth presentation is neither the first nor the last repetition of a given sequence). This analysis revealed that the majority of neurons in all layers exhibited repetition suppression (Fig. 2C). Statistical comparison revealed that the median number of suppressed neurons in the supragranular layers was significantly different from that of granular layer 4C (two-sample t-test, *α* = 0.05), while all other laminar comparisons were not significant. Lastly, we tested for statistical significance of repetition suppression on the single neuron level by comparing the observed responses to artificially created Poisson-distributed random spike trains (see Methods for details). We found that, across all laminar compartments, the majority of neurons showed a significant decrease in spiking, with only a handful of neurons showing significant spiking increases when the stimulus was repeated (Fig. 2D).

### Current Source Density Indicates Reduced Synaptic Activity

To characterize the origins of the laminar differences in repetition suppression described above, we computed Current Source Density (CSD), a quantitative measure of volume-averaged transsynaptic current flow (Mitzdorf 1985; Schroeder et al. 1998). In general, current sinks are linked to net depolarization while current sources are linked to passive return currents (Mitzdorf 1987; Schroeder et al. 1998).

The initial current sink in V1 granular layers that occurs following onset of visual stimulation is indicative of retinogeniculate synaptic activation (Mitzdorf 1987; Schroeder et al. 1998; Maier 2013; Godlove et al. 2014; van Kerkoerle et al. 2014). Subsequent current sinks in extragranular layers are believed to reflect net polarization due to subsequent, cortico-cortical processing (Douglas et al. 1988; van Kerkoerle et al. 2014; Cox et al. 2017). We thus wanted to determine the relative magnitude of the repetition-related CSD response reduction for each of these laminar compartments (Fig. 3A and Supplementary Fig. 6; see Supplementary Fig. 8 and 3 for the full sequence of LFP and spiking responses, respectively).

**Figure 3.**
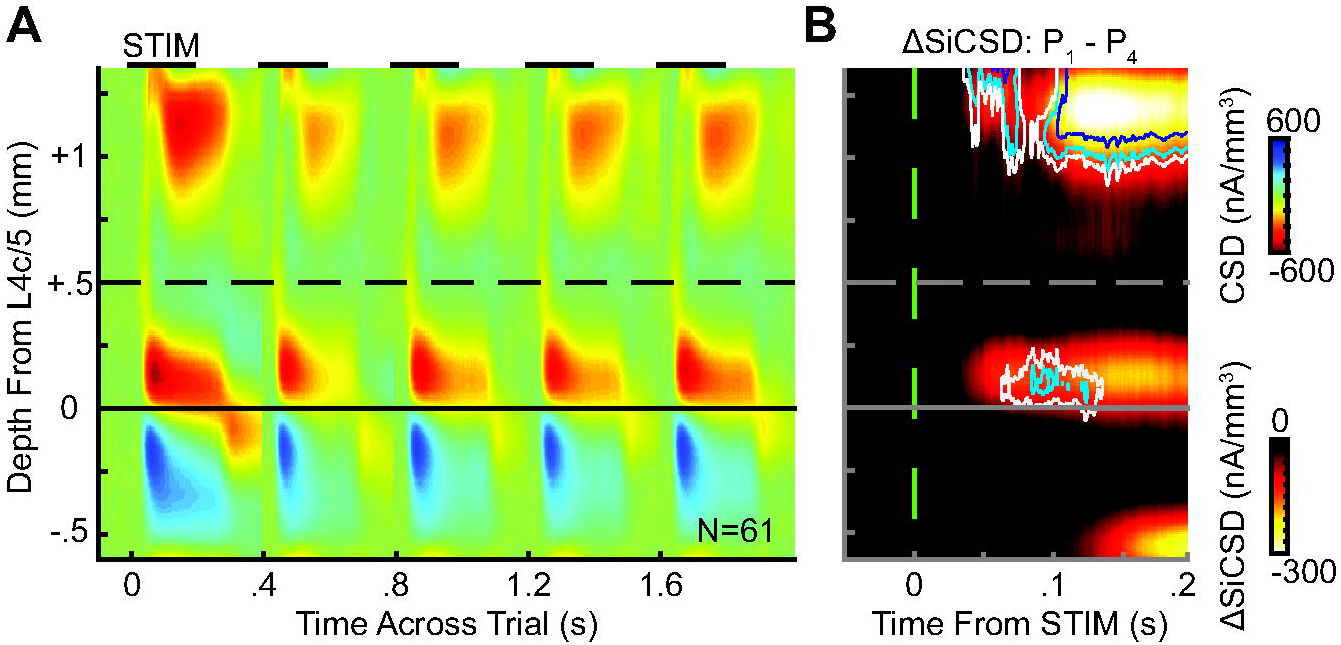
Repetition primarily modulates extragranular currents. (A) Mean CSD across presentations across trials. Stimulation denoted by bars at top. (B) Difference in CSD sinks (SiCSD) between first and fourth presentations, averaged across sessions. Interior of navy outline denotes *p* < 0.01, light blue *p* < 0.05, and white *p*< 0.1.

Inspection of the full CSD sequence reveals that the overall CSD response is largest for the initial stimulus presentation and smaller for all stimulus repetitions, suggesting that it repetition suppression is caused by a lack of synaptic activation rather than active inhibition. The smallest CSD response occurred for the second stimulus, coincident with the biggest differential in spiking (Fig. 3B), but CSD increased slightly for ensuing repetitions. The largest repetition-related decrease in current sinks was located in the supragranular layers, which harbor the bulk of cortico-cortical connections (Fig. 3B). This laminar differential in current flow suggests that repetition-related reduction in synaptic drive occurs outside the granular layer 4C as this layer receives only slightly (if at all) diminished reticulogeniculate input. Indeed, layer 4C CSD reduction did not reach significance until after significant supragranular response reduction took place (Fig. 3B, see also Supplementary Fig. 6). These findings suggest that the majority of repetition-related reduction in synaptic drive cannot be explained by adaptation of sensory input to V1. Instead, the bulk of suppression on the synaptic level arises from reduced intracortical signaling during the laminar propagation of sensory activation.

### Interocular Transfer of Response Suppression

Since the CSD analyses outlined above implicated a cortical origin of V1 repetition suppression, we tested this hypothesis by alternating stimulation between the two eyes. The two retinae receive no feedback from its projection targets in the primary visual pathway (Ortiz et al. 2017). And at the next step of visual processing, in the lateral geniculate nucleus (LGN), almost all neurons respond to one eye only (Zeater et al. 2015; Dougherty et al. 2018). Hence, sensory signals from the two eyes remain largely segregated before they arrive in V1. We thus reasoned that repetition suppression following interocular stimulus alternation would provide further evidence of an intracortical origin. In the same vein, if the repetition-related response reduction we observed was mostly due to sensory adaptation at earlier (monocular) stages of visual processing, V1 repetition suppression should be significantly reduced when successive stimulation alternated between the two eyes.

To perform this analysis, we divided our sample into units that were strongly biased towards one eye (“monocular” neurons) and units that responded to both eyes similarly (“binocular” neurons) (see Methods and Supplementary Fig. 7). We then compared the visual responses of each of these population – combined across all layers given the smaller sample size - across two conditions (Fig. 4): In the first condition, the initial stimulus was presented to the same eye as the repeating stimulus. In the second condition, the initial stimulus was shown to the dominant eye and the repeating stimulus was shown to the other eye. Our rationale for this analysis was that due to their monocular nature, neurons in the retina or the LGN would be insensitive to the stimulus repetition if adaptation occurred in the other eye. Only neurons that receive inputs from both eyes (binocular neurons) are expected to exhibit repetition suppression if the stimulus repetition affects the eye that was not initially stimulated.

**Figure 4.**
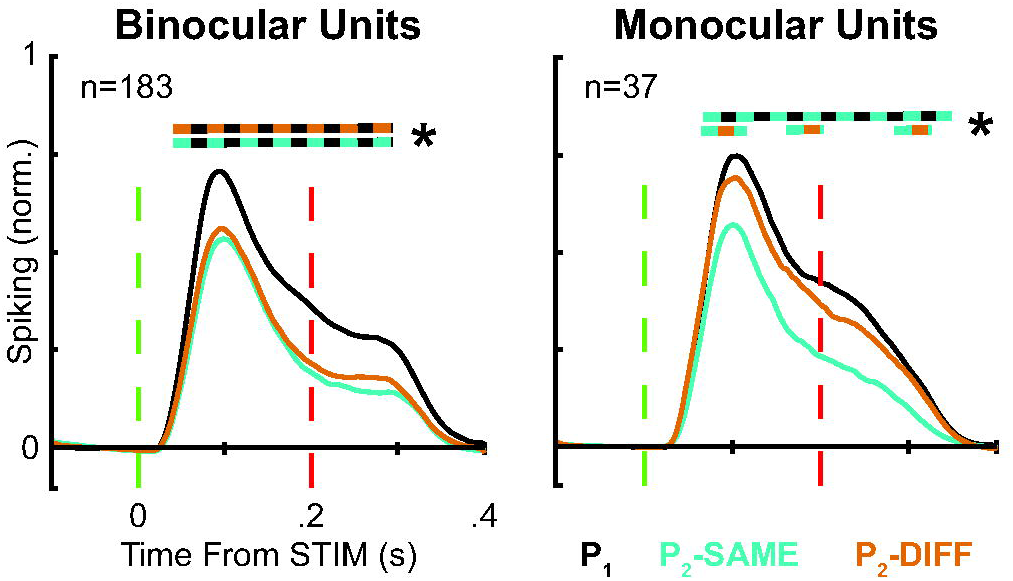
V1 repetition suppression in response to stimulating opposite eyes for binocular and monocular V1 neurons. Data represent the smoothed, median normalized response of all units that fall in each respective category. Black lines denote the response to initial presentation. In the case of monocular units, initial presentations were restricted to the dominant eye. Teal depicts the response to the second presentation if the initial presentation was presented to the same eye. Orange depicts the response to the second presentation when the initial presentation was in the other eye. Striped bars indicate times points that were significant at *p* < 0.05 (two sample t-tests).

Consistent with the notion of a cortical origin of repetition suppression, we found that monocular V1 neurons showed no significant reduction of activity when the initial stimulus was shown to one eye and the repeating stimulus to the other. In contrast, binocular V1 neurons, which constituted 84% of our sample, showed significant repetition suppression independently of ocular configuration (Fig. 4). In other words, the vast majority of V1 neurons showed repetition suppression independently of whether the same or opposite eyes underwent stimulus repetition. This finding suggests that V1 repetition suppression occurs even in the absence of sensory adaptation of monocular neurons in the retinae and the LGN further supports the idea of a cortical origin of repetition-related adaptation.

### Sharpened Orientation Tuning with Repeated Visual Stimulation

Having established that V1 repetition suppression is of cortical origin, we set to confirm earlier reports demonstrating enhanced encoding efficiency during these conditions. In particular, sharpening of neural tuning has been suggested as a mechanism to explain improved performance during repetitive stimulation (Desimone 1996; Wiggs and Martin 1998; Gotts et al. 2012). Indeed, sharpening of tuning has been shown to occur under a variety of stimulation conditions, including adaptation (Müller et al. 1999; Dragoi et al. 2001; Schoups et al. 2001; Felsen et al. 2002; Chen et al. 2005; Zinke et al. 2006; Hansen and Dragoi 2011; Buzsáki et al. 2012). This sharpening of tuning associated with repetition suppression is distinct from shifts in tuning preference that occur with more prolonged periods of exposure that lead to perceptual aftereffects (Dragoi et al. 2000; Dragoi et al. 2002; Felsen et al. 2002). We examined responses to various stimulus orientations during the initial presentation (Fig. 5A, black) and compared those to responses of repetitions (purple). This revealed that the response difference between the preferred and orthogonal orientations was greater for stimulus repetitions than for the initial presentation.

**Figure 5.**
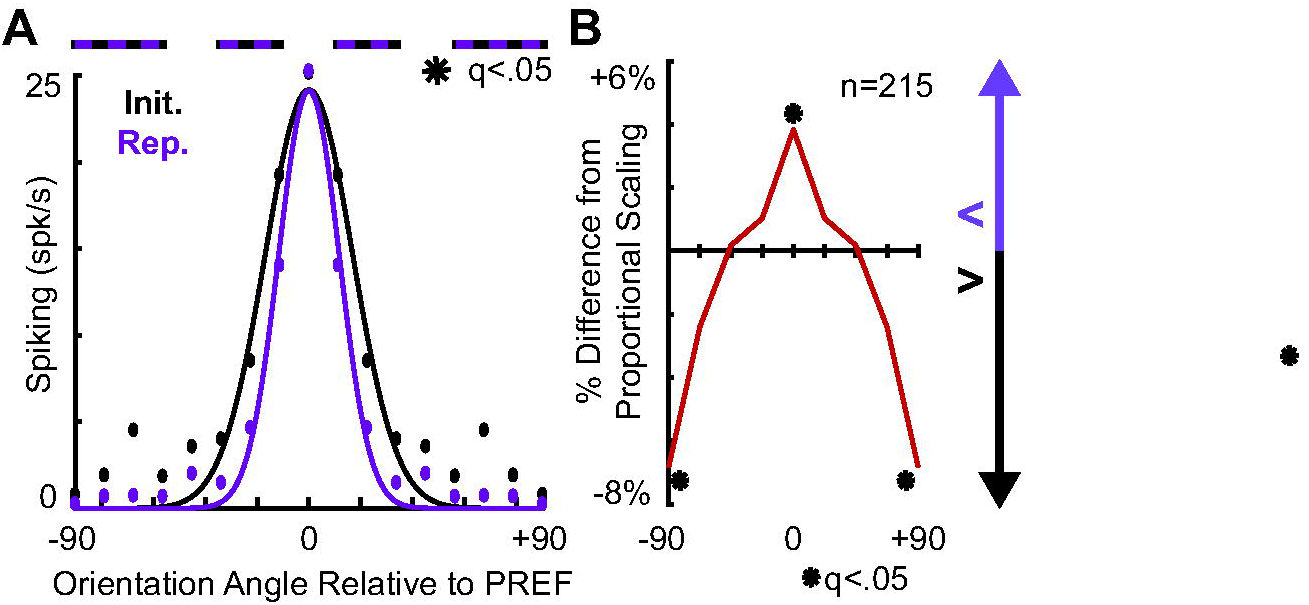
Sharpening of V1 orientation tuning with repetition. (A) Representative tuning curves from a single unit using a Gaussian fit (see Methods). Black represents responses from the initial presentation, purple, repetitions. Bars at top denote significant differences using a two-sample t-test with *p* < 0.05. (B) Population data showing deviation from proportional scaling. Asterisks denote values significantly different from 0 according to a t-test where *p* < 0.05 following FDR correction.

To quantify this effect, we calculated a tuning difference metric (see Methods). Of the 278 units we recorded, 215 showed significant orientation preference (*p* < 0.05, one-way ANOVA). The population average of tuned neurons revealed a significantly greater response than expected by proportional scaling for preferred orientations and a significantly reduced response at the orthogonal orientations during repetitions, resulting in a sharpened population tuning curve for the repetition-suppressed responses (Fig. 5B). This result is reminiscent of earlier reports (Hansen and Dragoi 2011) which found that orientation discrimination increases with stimulus adaptation.

### Stimulus Repetition Increases V1 Synchrony

As a final step, we investigated the effects of repetition on neural coherence. Neural synchrony can affect sensory processing independently of spike rates, and coherence has been shown to change with stimulation history (Gutnisky and Dragoi 2008; Hansen and Dragoi 2011). The spectral relationship between the local field potential (LFP) and the timing of spikes (spike-field coherence, or SFC) has been tied to neuronal synchronization within and between brain areas (Engel et al. 2001; Zeitler et al. 2006; Womelsdorf et al. 2007; Gilbert et al. 2010; Bosman et al. 2012; Spaak et al. 2012; Brunet et al. 2014; Schmiedt et al. 2014; Bastos et al. 2015; Dougherty et al. 2015; Haegens et al. 2015; Ninomiya et al. 2015). We thus sought to investigate whether SFC is enhanced in V1 when stimuli are presented repeatedly (see also Hansen and Dragoi 2011). To do so, we investigated the spectral content of LFPs recorded concurrently with spiking data to determine power in lower (theta, alpha, beta) and higher (gamma) frequency bands as a function of stimulus repetition (Supplementary Fig. 8). In line with earlier work (Gilbert et al. 2010; Hansen and Dragoi 2011), we found that stimulus repetition leads to a significant decrease of high-frequency (gamma) power concomitant with power increases in the low frequency (theta-beta) range (Fig. 6A). This pattern was consistent across the laminar compartments, which could be at least partially due to passive volume conduction (Katzner et al. 2009; Kajikawa and Schroeder 2011).

**Figure 6.**
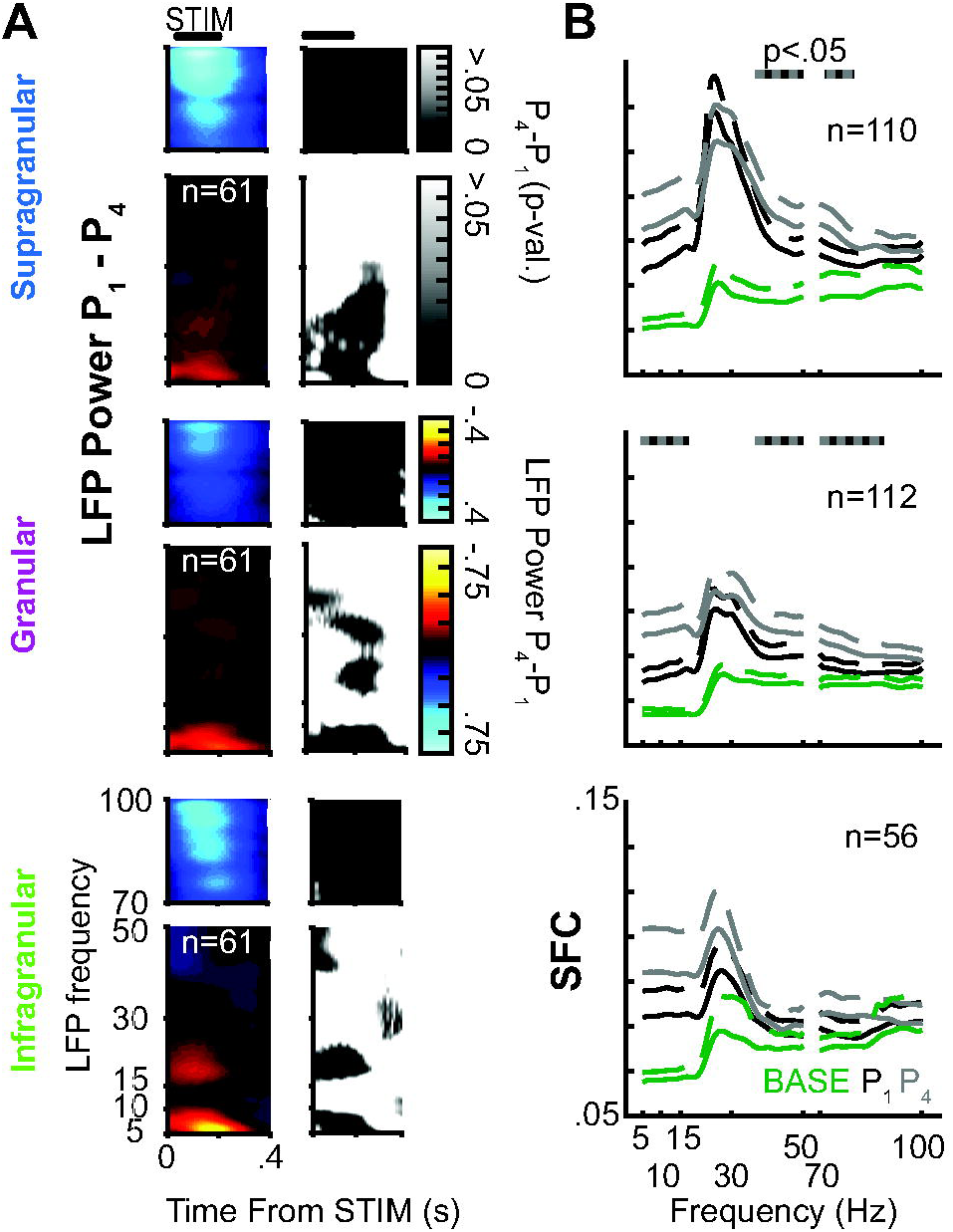
Changes in neural synchrony following stimulus repetition. (A) Comparison of LFP power during the initial and fourth presentation in normalized units (left) as well as p-values computed using a two-sample t-test (right) across 61 sessions. Reliable differences with repetition suppression are detected in low gamma, beta, and theta frequency bands across all laminar compartments. (B) Spike field coherence (SFC) following stimulus repetition. Solid lines denote mean SFC across units. Dashed line denotes the upper bound of the 95% confidence interval. Initial presentation (black) enhances broadband SFC relative to baseline (green). Stimulus repetition (gray) further enhances SFC across laminae. Gray bars on top denote statistical significance for the difference between initial presentation and stimulus repetition at *p* < 0.05 (two-sample t-test).

Since the above results suggests that LFP was modulated by stimulus repetition in a frequency-dependent manner, we next computed SFC for each condition to test for associated changes in neural synchrony. We found that SFC peaked in all layers across all conditions in the low beta range (~20-30Hz; Fig. 6B), reminiscent of earlier reports in macaque V1 (Hansen and Dragoi 2011) and V4 (Brunet et al. 2014). There was no shift in the spectral peak of the SFC functions between the initial stimulus and ensuing repetitions. However, SFC magnitude increased with successive stimulus presentations. Specifically, SFC in the alpha range was significantly enhanced in granular layer 4C and SFC in the gamma range was significantly elevated in both granular layer 4C and supragranular layers, suggesting that despite the decrease in spiking during repetitive stimulation, V1 (spike-field) coherence increased for neurons in the supragranular layers and granular layer 4C.

## Discussion

Our study is the first to our knowledge that demonstrates that stimulus repetition modulates the sequential pattern of sensory activation along the canonical cortical microcircuit. Through investigation of synaptic activation and binocular integration, we found that repeated stimulation evokes a similar initial response in the cortical input layers. It is only once this initial sensory activation propagates to the supragranular layers that adaptive changes become apparent, suggesting that stimulus repetition predominantly affects intracortical processing of sensory signals. This notion was further supported by our finding that V1 repetition suppression survives stimulus manipulations that reduce adaptation in the sensory periphery.

### Relation to Previous Findings

The reduced spike rates to repeated visual presentations observed in our study are reminiscent to the general component of repetition suppression while also mirroring earlier findings in early visual cortex of anesthetized cats (Movshon and Lennie 1979; Albrecht et al. 1984; Summerfield et al. 2008), rodents (King et al. 2016), and monkeys (Hansen and Dragoi 2011). The notion that V1 repetition suppression is predominantly of cortical origin is in line with earlier work in anesthetized macaques (Kohn 2007) as well as with recent optogenetic work in mice (King et al. 2016). We also replicated findings of earlier studies demonstrating sharpened neuronal tuning and heightened synchrony during stimulus repetition (Hansen and Dragoi 2011; Gotts et al. 2012; Brunet et al. 2014), suggesting that both of these mechanisms coincide with the adaptation of intracortical signaling (Fig. 7). The resulting enhanced efficiency of sensory encoding upon stimulus repetition might underpin the improved perceptual performance given a familiar environment, while reducing metabolic cost (Sengupta et al. 2013).

**Figure 7.**
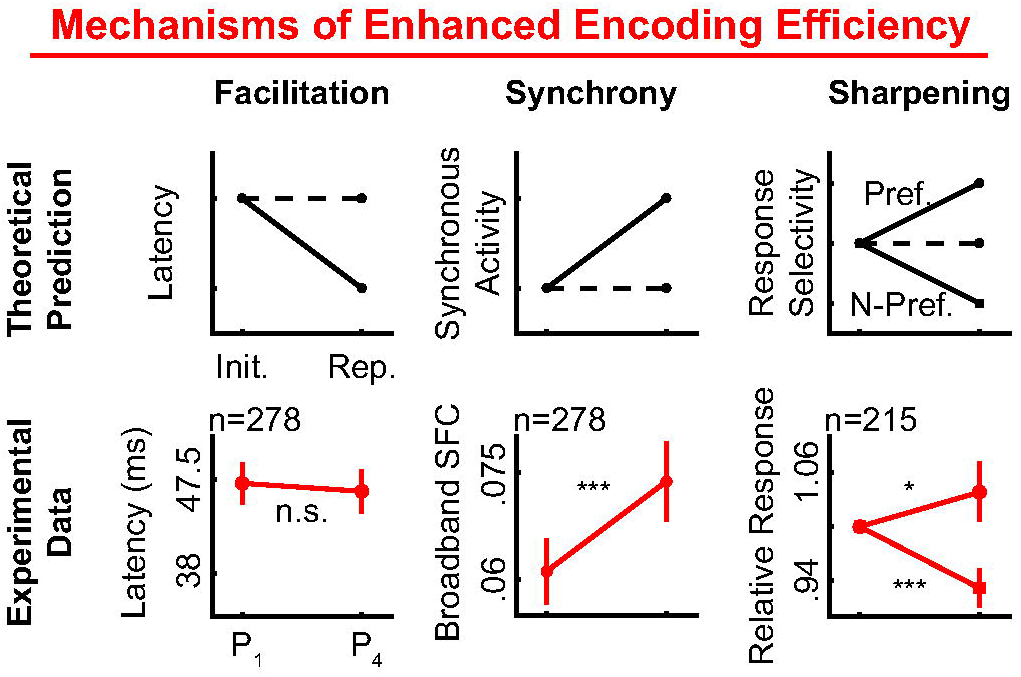
Enhanced stimulus encoding concurrent with suppression of visual responses. Three models of enhanced encoding efficiency with repetition suppression were tested in this study. Predictions outlined in the first row. Solid lines depict the trend predicted by the respective model. Null effects depicted by dashed lines. Data are shown in red. Statistics were performed using one- or two-sample t-tests comparing the response to the first and fourth presentations of each sequence. No evidence was found for the facilitation model (i.e. reduced response latency; *p* > 0.05). However, predictions of synchrony and sharpening models reached significance in the form of increased SFC and deviation from proportional scaling (\**p* < 0.05, \*\*\**p* < 0.001), respectively.

### Putative Mechanisms

The greatest response difference caused by stimulus repetition was a diminished synaptic response in the supragranular layers that harbor the bulk of cortico-cortical connections (Fig. 8). Why the temporal profile of CSD deviates from that of single-unit spiking is unclear and will be the focus of upcoming work. Previous studies found that repetitive stimulation to the same retina causes adaptation (Victor 1987; Berry et al. 1999; Baccus and Meister 2002). Assuming such adaptation to repeated stimulation in the retina and the LGN, we can conclude that this cortico-cortical reduction either occurred independently of - or amplified - a much smaller decrement in initial sensory activation, as the difference in initial synaptic activation in granular layer 4C was too small to reach significance in our sample. This interpretation is consistent with earlier work in anesthetized macaques and rodents, showing weaker adaptation in LGN compared to V1 (Kohn 2007; King et al. 2016). Anatomically, it is also possible that LGN neurons inherit some aspects of repetition suppression via corticofugal feedback from V1. However, optogenetic inactivation of rodent V1 does not abolish sensory adaptation in the LGN (King et al. 2016). Whether the origin of the reduced cortico-cortical response lies within V1 (e.g., via diminished lateral connections) or is derived from downstream stages in the visuocortical hierarchy (e.g., via diminished feedback to V1) cannot be determined from the laminar pattern of CSD alone.

**Figure 8.**
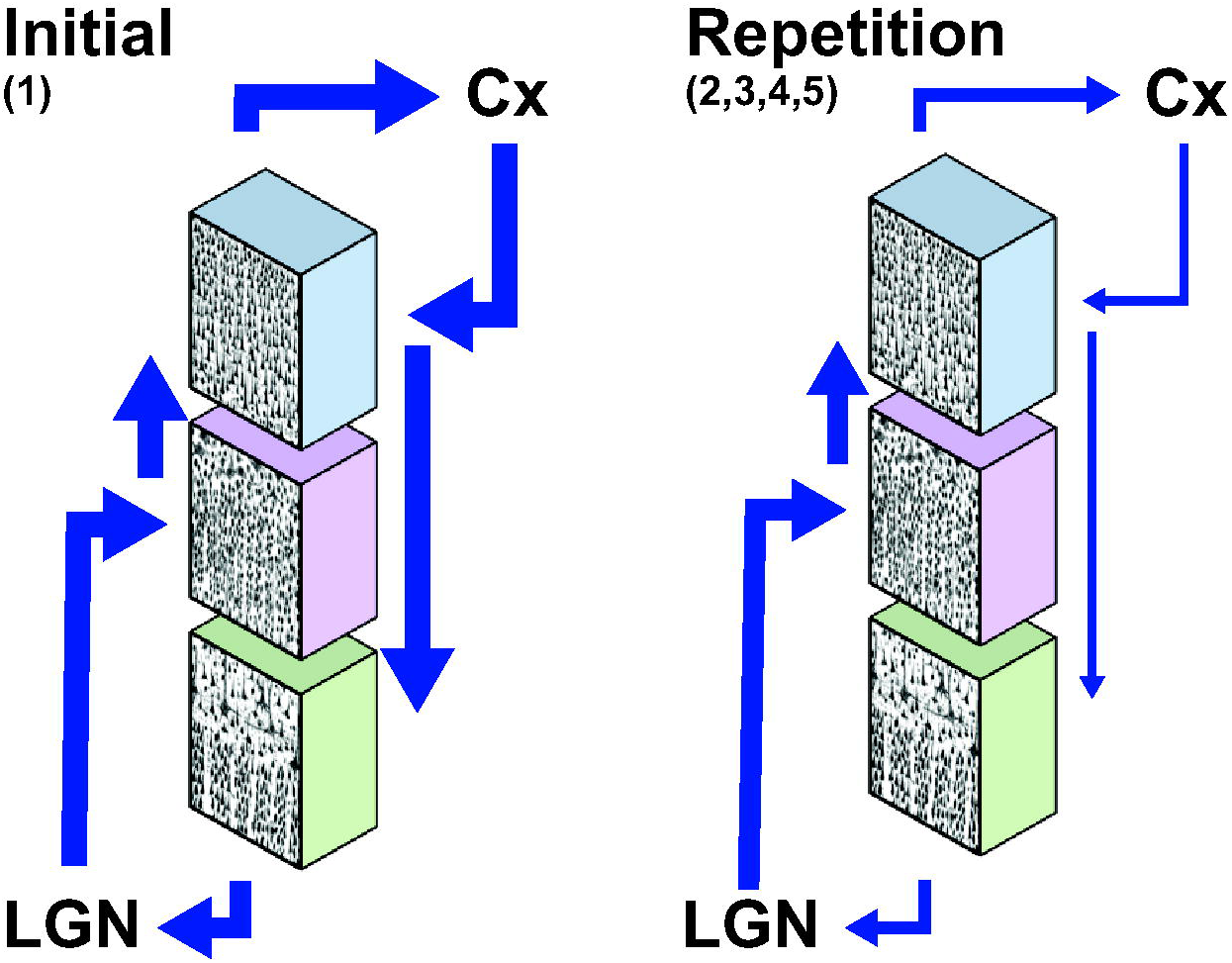
Cartoonized model of functional adaptation of V1’s laminar microcircuit. When a stimulus is first encountered, sensory activation initially mainly activates spiny stellate cells in the granular input layer 4C of V1 (pink). Layer 4C neurons predominantly project onto supragranular neurons (blue), whose activation spreads to other layers (green) and eventually to other visual areas downstream the visual hierarchy. Intracortical spread of activation via horizontal connections as well as feedback from downstream areas to the extragranular layers of cortex can further enhance neuronal responses in these layers (left). Following stimulus repetition (right), intracortical activation is reduced, leading to reduced overall activity.

### V1 Suppression without Increased Inhibition

Another important conclusion derived from our finding of decreased synaptic activity in extragranular layers when stimuli are repeated is that intracortical signaling is reduced under these conditions. This observation suggests that while intracortical processing plays a profound role for adaptation, the absence of intracortical activation underlies most repetition suppression in V1. In other words, repetition suppression in V1 does not seem to be caused by active inhibition from increased feedback from higher level areas, as heightened inhibition or increased feedback should be reflected in increased synaptic activation and thus a larger CSD response compared to the initial presentation of the stimulus (Spaak et al. 2012; van Kerkoerle et al. 2014; Cox et al. 2017; van Kerkoerle et al. 2017). Having said this, it should be noted that CSD alone cannot provide conclusive evidence regarding the precise synaptic mechanisms underlying repetition suppression since it represents a spatially summed population signal. Identification and experimental interrogation of inhibitory interneurons under a repetition suppression paradigm would allow for more direct testing of this hypothesis.

### Relationship to Repetition Suppression in Extrastriate Cortex

The phenomenon described here both resembles as well as differs from reports of repetition suppression in IT cortex (Miller et al. 1991, 1993; Miller and Desimone 1994; McMahon and Olson 2007). Both IT as well as V1 responses exhibit the greatest suppressive effect following the initial stimulus. However, IT neurons continue to adapt (i.e., further diminish their responses) with repeated presentations, while our V1 neurons maintained a more or less stable diminished response following the third stimulus repetition. Stimulus differences and other parameters make it difficult to directly compare results obtained in these separate visual areas. However, the fact that a similar adaptive process is seen at both the initial cortical step of visual processing as well as near the apex of the visual cortical hierarchy raises the question whether all areas that fall in between exhibit a similar profile of adaptation (see also Brunet et al. 2014; Patterson et al. 2014).

### Potential Role of Expectation

Repetition suppression has been linked to predictive coding and expectation more generally (Summerfield et al. 2008; Auksztulewicz and Friston 2016). In this framework, reduced spiking responses to repeated stimuli are due to the reduction of a prediction error signal when probabilistic expectations about stimulus reoccurrence are met. We have not directly tested this hypothesis, which can be done by altering the statistical distributions of stimulus occurrence, for example. Single neurons recordings in the IT using such manipulations did not yield conclusive results (Kovács and Vogels 2014; Bell et al. 2016). Moreover, we would expect our paradigm to evoke a certain prediction error despite the repetitive stimulation given that at least one stimulus feature (orientation or ocularity) was randomly varied between presentation (although we held all other variables such spatial frequency, contrast level and stimulus size constant). It is noteworthy, however, that our data seem compatible with a recently proposed model of predictive coding along V1’s columnar microcircuit (Bosman et al. 2012) in that we observed the model-predicted reduced synaptic activation in V1’s feedback-recipient layers for repeated stimuli. Note that we deliberately limited our study to the effects of stimulus repetition on sensory cortical activation. Previous studies have shown that spontaneous activity (i.e., ongoing spiking in between stimulus presentations) in sensory areas can be affected by adaptation and memory of the most recent sensory stimulus as well as anticipation (Bisley and Pasternak 2000; van Kerkoerle et al. 2017). Visual inspection of both the population spiking and LFP responses from our paradigm suggests that spontaneous activity during the inter-stimulus intervals was also modulated by the task. This finding may also be related to the possibility of “neuronal entrainment” (Lakatos et al. 2008; Schroeder and Lakatos 2009; Morillon et al. 2016) to the rhythmic stimulation sequence (which is challenging to avoid without adding the additional confound of varying time constants), but we will investigate this phenomenon in a separate study.

### Role of Attention for Repetition-Related Response Reduction

One alternative explanation to the observed reduction in spiking could be that the animals’ attention fades over time across each stimulus sequence. Indeed, it is possible that the initial visual stimulation of each sequence draws exogenous attention towards the stimulus location, resulting in increased neuronal firing and enhanced perceptual sensitivity. With subsequent stimuli, this heightened state of attention might fade or even be inhibited (i.e., by inhibition of return). As a result, neuronal responses would be expected to decline and, importantly, behavioral performance would decrease accordingly. As we did not control the animal’s state of attention, we cannot entirely rule out this hypothesis. However, the fact that behavioral performance generally increases rather than decreases with stimulus repetition (Tulving and Schacter 1990) suggests that attention does not play a significant role in repetition suppression and the associated effects we observed.

## Authors’ Contributions

J.A.W. and A.M. conceptualized the study. J.A.W., M.A.C., and K.D. collected data. J.A.W. and M.A.C. preprocessed the data. J.A.W. performed analysis and prepared visualizations. J.A.W. and A.M wrote the original draft, and all authors contributed to revisions.

## Funding

This work was supported the National Eye Institute at the National Institutes of Health (grant numbers 1R01EY027402-01 and 5T32EY007135-23 to J.A.W. and K.D.).

## Acknowledgements

The authors would like to thank B. and R. Williams, M. Schall, E. A. Sigworth, M. R. Feurtado, and Dr. W. Zinke for technical support as well as Dr. J. D. Schall and K. A. Lowe for comments on an earlier version of the manuscript. *Conflict of interest: None declared*.

